# Tumour suppressors LKB1 and SMARCA4 functionally interact to regulate gene expression of diverse biological processes in lung cancer

**DOI:** 10.64898/2026.01.26.699134

**Authors:** M. Bourouh, JH. Kim, PA Marignani

## Abstract

The tumour suppressor kinase LKB1 is known to regulate the activity of the metabolic sensor AMPK, that when under energy stress shifts metabolism from anabolism to catabolism, thus linking LKB1 to AMPK-mediated gene expression. Coupled with its role as a tumour suppressor kinase, LKB1 is an important metabolic regulator implicated in multiple malignancies and is frequently mutated in lung cancer. Previously, we discovered that LKB1 binds to the SWI/SNF chromatin remodelling ATP-dependent helicase subunit SMARCA4, directly linking LKB1 to gene expression. How LKB1 and SMARCA4 collaborate to regulate gene expression in lung cancer has not been well characterized. We used an *in-silico* approach to explore how LKB1 and SMARCA4 may cooperate to regulate gene expression. We mined our previous scRNA-seq dataset from 4 lung cancer cell lines with differential *LKB1* and *SMARCA4* expression status to identify genes regulated by both LKB1 and SMARCA4. We correlated our results using bulk RNA-seq results from human lung tumours. We show that LKB1 and SMARCA4 likely function together to regulate gene expression in multiple biological processes in lung cancer cell lines. Gene expression profiles from LKB1 and SMARCA4 mutant cells are similar, suggesting LKB1 and SMARCA4 function in a linear pathway to regulate gene expression. Furthermore, we observed similar results in human lung tumours, particularly in late-stage disease. We propose a model where LKB1 acts as a nexus between metabolism and gene expression, acting via the SMARCA4-SWI/SNF complex to regulate gene expression in lung cancer.

## Introduction

Metabolic reprogramming plays a crucial role in cancer cells’ ability to thrive in a poor nutrient environment and regulate gene expression through metabolic signalling intermediates. Epigenetic enzymes utilize key metabolic intermediates such as ATP, acetyl-CoA, and S-adenosylmethionine (SAM) to modulate gene expression through histone modifications (1–3). Epigenetic histone modifications alter the chromatin landscape, generating landmarks at gene regulatory elements to differentiating transcriptionally active chromatin regions from transcriptionally silent. The switch/sucrose non-fermenting (SWI/SNF) chromatin remodelling complexes for example hydrolyses ATP to displace nucleosomes, mobilize histones, and open chromatin to regulate gene expression (4, 5). The availability of ATP, acetyl-CoA and SAM can drastically alter the global chromatin landscape, and the availability of these cofactors are determined by the metabolic state of cells. This mechanism allows cells to leverage metabolic output to influence cellular responses through epigenetic regulation of gene expression. Mutations in tumour suppressor genes that disrupt metabolic pathways can lead to aberrant epigenetic changes commonly associated with malignant transformation.

Important regulators of metabolism and epigenetics are tumour suppressors Liver kinase B1 (LKB1) and SWI/SNF related, matrix associated, actin dependent regulator of chromatin, subfamily a, member 4 (SMARCA4), that are often mutated or lost in non-small cell lung cancer (NSCLC) (6, 7). Interestingly, *LKB1* and *SMARCA4* are also located in close proximity on chromosome 19p13.3, and loss of heterozygosity (LOH) is frequently observed (6, 7). The best characterized role of LKB1 is in energy metabolism, where LKB1 activates AMP-activated protein kinase (AMPK) to shift metabolic pathways from anabolism to catabolism (8–10). SMARCA4 is the ATP-dependent helicase subunit of the SWI/SNF chromatin remodelling complex, hydrolysing ATP to displace nucleosomes, opening chromatin for transcription factors to bind gene regulatory elements (4, 5). Both *SMARCA4* and *LKB1* mutant NSCLC cases are associated with smokers and consequently *KRAS* mutations. *LKB1* is mutated in approximately 20% of NSCLC cases (11, 12), while*SMARCA4* is mutated in 8% of NSCLC cases, with 39% of those cases representing *LKB1-SMARCA4* co-mutation (13, 14).

Our lab previously discovered that nuclear localized LKB1 binds to SMARCA4 *in vivo*, specifically, LKB1 binds to the helicase domain of SMARCA4, promoting SMARCA4 ATPase activity, independent of LKB1 catalytic activity (15). ATPase experiments *in vitro*, showed that in the presence of DNA, the ATPase activity of SMARCA4 is 3-fold higher in the presence of LKB1 compared with the presence of SL26, a catalytic deficient mutant of LKB1 first identified in Peutz-Jeghers syndrome (16). This strongly suggests that the binding of SMARCA4 to LKB1 is necessary for the ATPase activity of SMARCA4, and that the ATP-dependent chromatin remodelling function of SMARCA4 is reliant on LKB1 (15). We have also shown that the tumour suppressor function of SMARCA4 is partially dependent on LKB1. When SMARCA4 is expressed in SW13 cells, a SMARCA4 mutant cell line that expresses endogenous LKB1, SW13 cells undergo cell cycle arrest. The cell cycle arrest is suppressed when SMARCA4 is co-expressed with SL26 but not LKB1, indicating the catalytic activity of LKB1 is required for SMARCA4 mediated cell cycle arrest (15). In a later study, we discovered that LKB1 catalytically deficient mutants promote the expression of *CYCD1* by directly binding the *CYCD1* promoter (17). Since SMARCA4 promotes the expression of *CDKN1A* (P21), leading to the inhibition of CDK4-CYCD1 that reduces the phosphorylation of retinoblastoma (RB) thus repressing E2F transcription factors, ultimately inhibiting the expression of G1/S factors (18). More recently, the loss of binding between LKB1 to SMARCA4 leads to PRC2-dependent transcriptional inhibition through increased H3K27me3. This leads to upregulation of oxidative stress pathways, and amino acid metabolism dysregulation (19).

Indirect observations implicate LKB1 and SMARCA4 in lipid metabolism, a well characterized function of LKB1. The LKB1-AMPK pathway can regulate expression of genes involved in fatty acid (FA) biosynthesis. Phosphorylation of sterol regulatory element-binding protein 1 (SREBP1), a transcriptional co-activator, by AMPK results in inhibition of SREBP1, resulting in deregulation of FA biosynthesis genes (20, 21). SMARCA4 can bind to SREBP1, suggesting that SMARCA4 plays a role in regulating expression of FA biosynthesis genes, indirectly linking SMARCA4 to LKB1 functions (22). In addition to SREBP1, the LKB1-AMPK signalling pathway is involved in regulating β-oxidation through peroxisome proliferator-activated receptors (PPAR) nuclear receptors. Here, LKB1-AMPK signalling activates PPARα in skeletal muscle cells, which leads to increased transcription of gene involved in β-oxidation (23, 24). SWI/SNF complexes also play a role in PPAR pathway, as the SWI/SNF complex is recruited to PPARγ target genes (25). These results suggest indirectly that the interaction between LKB1 and SMARCA4 regulates lipid metabolism.

Finally, LKB1 and SMARCA4 both regulate transcription mediated by the ERα receptor (26, 27). LKB1 binds to ERα independent of catalytic activity, albeit LKB1 catalytic activity is required for transcription of ERα target genes. Interestingly, ERα can recruit SMARCA4 to ERα target genes, where this is dependent on histone acetylation. Here, LKB1 and SMARCA4 may function together to regulate expression of ERα target genes, and histone modifications play an important role (26, 27).

LKB1 has also been linked to chromatin organization independent of SMARCA4 (2, 28). One study examined the effect LKB1 has on chromatin accessibility using assay for transposase accessible chromatin using sequencing (ATAC-seq) after restoring *Lkb1* expression in *Lkb1* mutant mouse lung tumour cells and found more than 30,000 genomic loci exhibited changes in chromatin accessibility (28). Furthermore, another study linked *Lkb1* loss in mouse lung tumours to upregulation of SAM production, leading to global histone methylation and repression of retrotransposons (2).

The consequences of *Lkb1* chromatin-remodelling function have been observed in collaboration with *Kras* mutant lung cancer in mice models. It was observed that *Lkb1* loss in *Kras* lung cancer mouse models exhibit loss of H3K27me3 and gain of H3K27ac and H3K4me3 histone modifications. These modifications correlated with transition from lung adenocarcinoma subtype to squamous cell carcinoma, indicating that this epigenetic change was facilitated through *Lkb1* loss and suggested that *Lkb1* loss impacted differential subtypes of lung cancer progression mediated by histone modifications (29).

Independent of Kras, Lkb1 functions to regulate chromatin dynamics in pancreatic β cells. *Lkb1* loss was correlated with dysregulation of FOXA, MAFA, and RFX6 transcriptional regulatory elements. This highlights a specific role of LKB1 in regulating chromatin dynamics in a variety of cells independent of KRAS and SMARCA4 (30).

These studies highlight the functional role of LKB1 in regulating chromatin remodelling via SMARCA4, however, the involvement of SMARCA4 in LKB1-dependent pathways and gene expression regulation has not been well characterized. To elucidate the functional relationship between LKB1 and SMARCA4 in lung cancer, we conducted scRNA-seq on four lung cancer cell lines with varying *LKB1* and *SMARCA4* mutation status (31, 32). Our analysis revealed that LKB1 and SMARCA4 function together in a linear pathway to modulate overlapping gene expression profiles in lung cancer cells. Notably, we observed that the expression profiles associated with *LKB1* and *SMARCA4* become apparent in late-stage human lung tumours. Our findings support a model wherein LKB1 functions dually as both a metabolic regulator, acting through AMPK signalling, and as a transcriptional regulator, acting through the SMARCA4-SWI/SNF chromatin remodelling complex. Consequently, our model positions LKB1 as a crucial nexus bridging metabolic processes and gene expression regulation in lung cancer.

## Results

### LKB1 and SMARCA4 regulate expression of overlapping genes

To characterize the interaction between LKB1 and SMARCA4 we mined our scRNA-seq dataset performed on 4 well-characterized lung cancer cell lines with varying *LKB1* and *SMARCA4* expression status; Calu-3, H460, H1299, and A549 (**Figure 1A**) (31, 32). Calu-3 cells (WT) are wild-type for both *LKB1* and *SMARCA4*, therefore will be referred to as WT. H1299 cells (*S*) are *SMARCA4* deficient, H460 cells (*L*) are *LKB1* deficient, and A549 cells (*LS*) are *LKB1, SMARCA4* deficient (**Figure 1A**) (33). All of these mutations do not produce functional protein (31, 33, 34). To classify differentially expressed genes (DEGs), we compared log_2_ fold change (log_2_FC) of all 11002 DEGs detected in our scRNA-seq dataset between *L*vsWT, *S*vsWT, and *LS*vsWT (**Table S1**). We classified DEGs as either LKB1-specific (L-DEGs), SMARCA4-specific (S-DEGs), or LKB1-SMARCA4-specific (LS-DEGs) based on the DEG expression profile. We reasoned that L-DEGs are detected only when *LKB1* is mutated (*L*vsWT and *LS*vsWT, n=911) and S-DEGs are only detected when *SMARCA4* is mutated (*S*vsWT and *LS*vsWT, n=814) (**Figure 1B**). LS-DEGs comprise the largest classification (n=5528, ∼50% of genes detected) and represent DEGs detected when either *LKB1* or *SMARCA4* are mutated (*L*vsWT, *S*vsWT, and *LS*vsWT) (**Figure 1B**). Therefore, our results suggest LKB1 and SMARCA4 are important regulators of gene expression and may largely function together to regulate expression of common genes.

**Figure 1.**
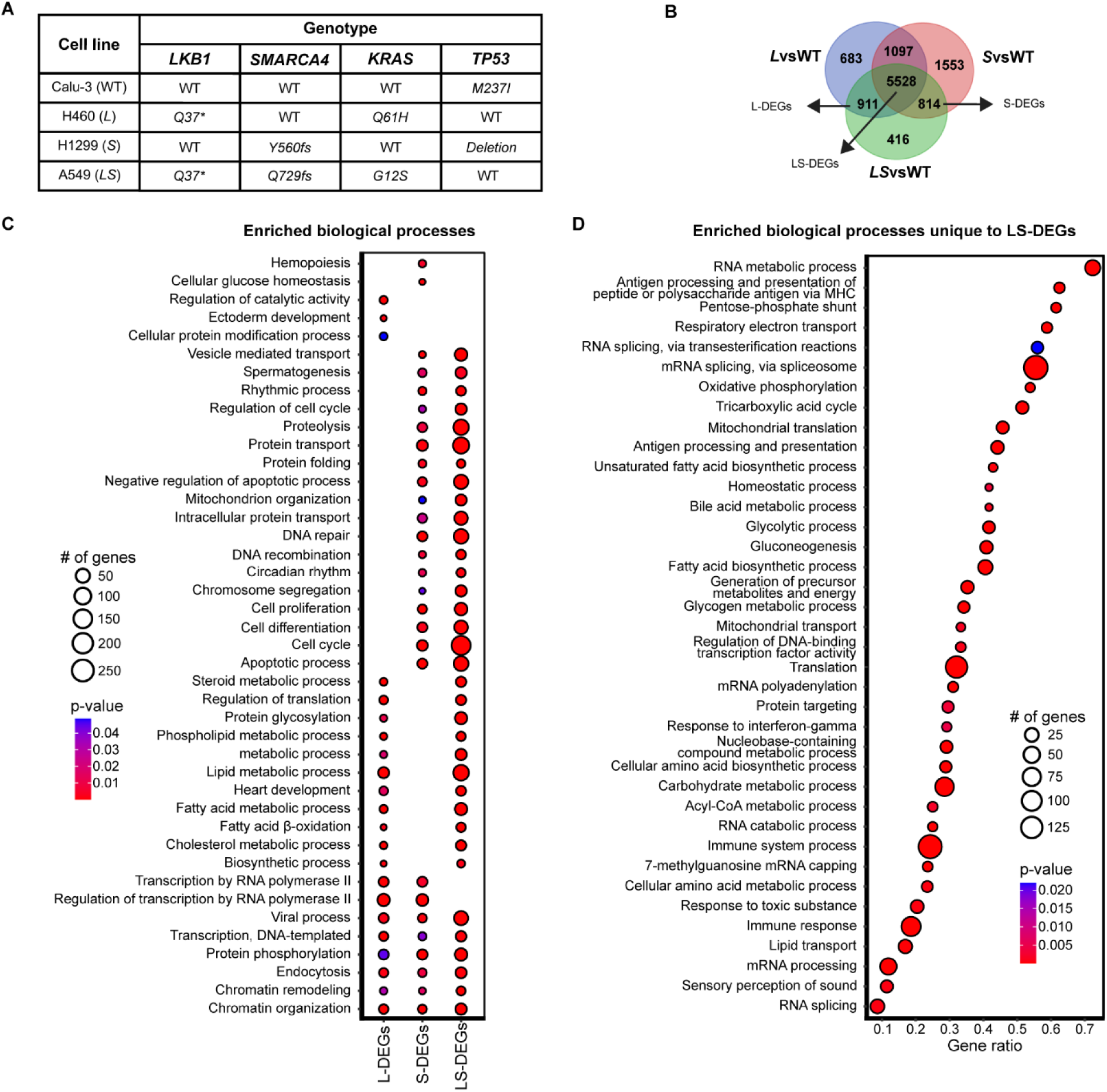
scRNA-seq on lung cancer cell lines identifies LKB1 and SMARCA4 gene expression signature. **A**) Genotype of lung cancer cell lines (Calu-3, H460, H1299, A549) used for scRNA-seq. **B**) Venn diagram representing the classification of DEGs L-DEGs, S-DEGs and LS-DEGs from comparing *L*vsWT, *S*vsWT, and *LS*vsWT DEGs. Number of DEGs classified as L-DEGs (n=911), S-DEGs (n=814), and LS-DEGs (n=5528). **C, D**) Dot plot of enriched biological process (BP) terms from GSEA analysis. Enriched BP terms for L-DEGs, S-DEGs, and LS-DEGs (**C**) and unique to LS-DEGs (**B**). Gene ratio in **D** calculated by number of genes/term size. Dot size represents the number of genes identified in each BP term, colour represents p-value with red representing low p-value and blue represents high p-value. Only BP terms that are statistically significant (*P*<0.05) and have >3 genes are shown.

### LKB1 and SMARCA4 have considerable overlapping functions

We then performed gene set enrichment analysis (GSEA) on L-DEGs, S-DEGs, and LS-DEGs to identify overrepresented gene ontology (GO) terms in the category of biological process (BP) - (See materials and methods) (**Table S2**). L-DEGs were enriched in biological processes related to metabolic pathways, particularly in fatty acid and cholesterol metabolic processes, consistent with well characterized LKB1 functions (**Figure 1C**) (8, 17, 35, 36). S-DEGs were enriched in diverse biological processes including protein transport, cell cycle, cell proliferation, and DNA repair (**Figure 1C**). L-DEGs and S-DEGs displayed overlapping enrichment in biological processes related to gene expression, depicted by overlapping functions in transcription, chromatin organization and chromatin remodelling (**Figure 1C**).

LS-DEGs represented the overwhelming majority of classified DEGs. If these DEGs represented overlapping functions of LKB1 and SMARCA4, LS-DEGs would be enriched for similar BP terms as L-DEGs and S-DEGs. Enriched BP terms for LS-DEGs showed significant overlap with enriched biological process of L-DEGs and S-DEGs, suggesting that LS-DEGs were enriched for genes that represent LKB1 and SMARCA4 functions (**Figure 1C**) In addition, LS-DEGs also exhibited unique enrichment in various biological processes implicated by LKB1 and SMARCA4 (**Figure 1D**). Many overrepresented BP terms corresponded to metabolic processes, but we also observed enrichment in translation, transcription, immunity and cell cycle biological processes (**Figure S1A**). Therefore, our workflow identified both unique and overlapping functions of LKB1 and SMARCA4 consistent with previously characterized LKB1 and SMARCA4 functions (8, 17, 35).

### *LKB1* and *SMARCA4* mutant human tumours exhibit similar expression profiles

The largely overlapping BP terms suggest that LKB1 and SMARCA4 function together to regulate gene expression of common pathways using cells in culture. To explore the translational consequences of this interaction in lung cancer, we asked whether our scRNA-seq results could be recapitulated using lung tumour transcriptional data from the cBioPortal cancer genomics database (37–39). It has been shown that *KRAS-LKB1* lung tumours exhibit different transcriptional and phenotypic properties than *KRAS-TP53* tumours,(40) however, the comparison between *LKB1* and *SMARCA4* tumours has not been investigated. We examined bulk RNA-seq data of tumours from the cBioPortal database, using tumours that had similar genetic background as the cell lines used for our scRNA-seq analysis; Calu-3, H460, H1299 and A549, with *TP53* mutant tumours representing Calu-3 (**Figure 1A** and **Table S3**). We first explored the overall transcriptional similarity between cell lines and primary lung tumours. We performed a correlation analysis to examine transcriptomic relationships among the 4 tumour types *TP53*, *LKB1*, *SMARCA4*, and *LKB1, SMARCA4* double mutant, and compared the correlation matrices of the tumours to the cell lines used in our scRNA-seq data. We observed transcriptional similarity between cell lines and tumours mutant for *LKB1* (H460, A549) and there is transcriptional similarity between *LKB1* and *LKB1-SMARCA4* tumours (**Figure 2A**). Moreover, *SMARCA4* and *TP53* tumours expression profiles are correlated, like the comparison of H1299 (*SMARCA4* mutant) and Calu-3 cells (*TP53* mutant) (**Figure 2A**). These results revealed that there is transcriptional similarity between the cell lines used in our scRNA-seq and cBioPortal primary lung tumour bulk-RNA-seq datasets.

**Figure 2.**
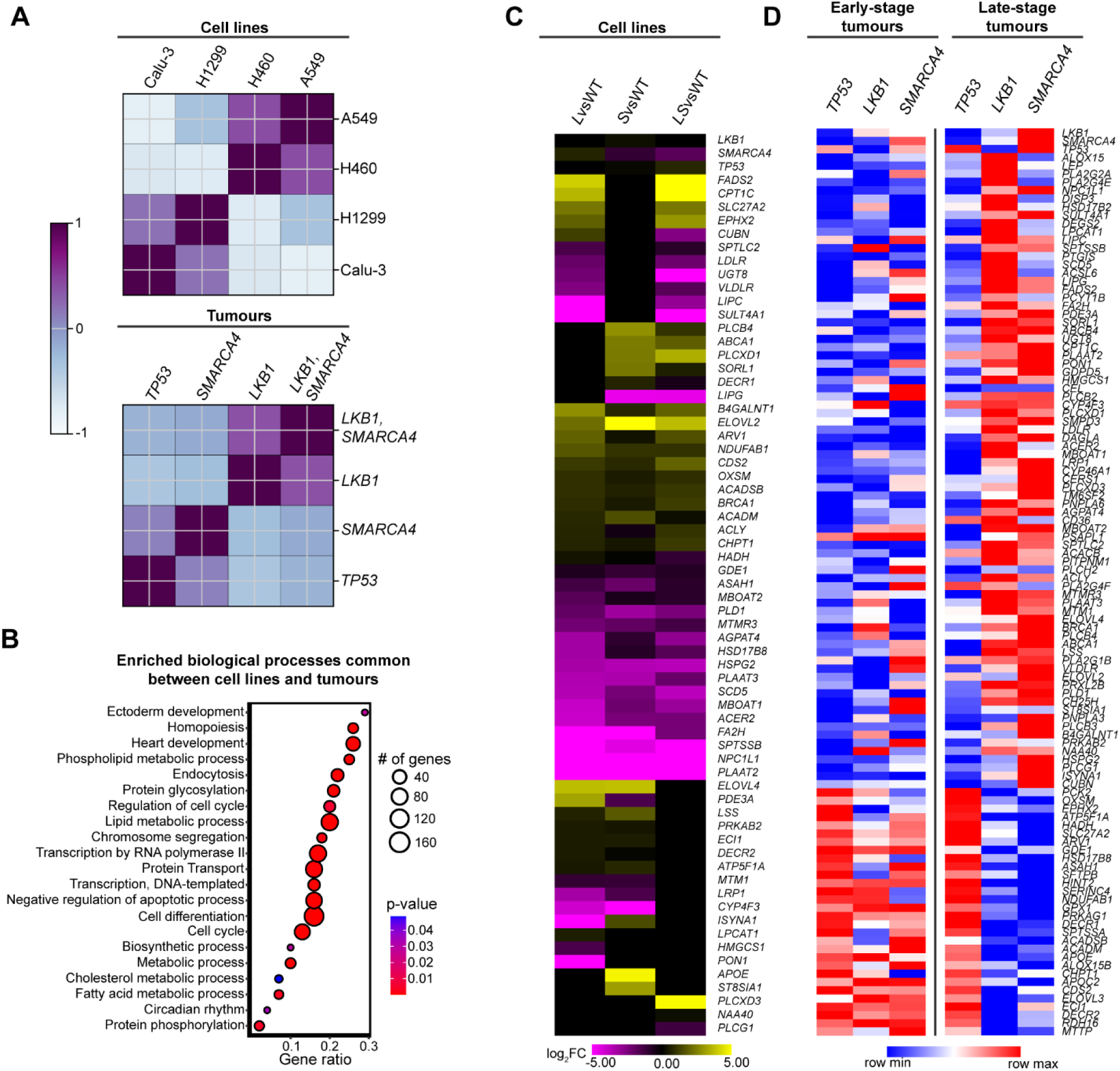
LKB1 and SMARCA4 Lung tumours exhibit similar expression profiles. **A**) Correlation analysis between scRNA-seq of lung cancer cell lines Calu-3, H460, H1299, and A549 and *TP53*, *LKB1*, *SMARCA4*, and *LKB1,SMARCA4* human lung tumours from cBioPortal database. Cell line or genotype of tumour is listed on the top and right of two pair-wise matrix plots. Colour represents z-score of the pair-wise matrix plots. Blue represents low z-score (low correlation) while purple represents high z-scores (high correlation). **B**) Dot plots of enriched BP terms common between cell lines and tumours (L-DEGs, S-DEGs, LS-DEGs and *LKB1* and *SMARCA4* mutant tumours). Gene ratio represents the number of genes/term size. Dot size represents the number of genes identified in the term, colour represents p-value with red representing low p-value and blue represents high p-value. Only BP terms that are statistically significant (*P*<0.05) and have >3 genes are shown. **C, D**) Heatmap plot of expression of DEGs identified from GSEA analysis from *LKB1* and *SMARCA4* early-stage and late-stage tumours involved in lipid metabolic process. **c**) The heatmap plot represents expression of genes from *L*vsWT, *S*vsWT, and *LS*vsWT using scRNA-seq data (purple/yellow). Yellow represents high expression, and purple represents low expression with scale of −5 to 5 log_2_FC. DEGs not detected or have no change in expression are represented in black. **D**) The heatmap plot represents expression of genes from *TP53*, *LKB1*, and *SMARCA4* early-stage and late-stage tumours (red/blue). Red represents high expression, and blue represents low expression with scale of row min row max.

To investigate pathways implicated in *LKB1* and *SMARCA4* tumours, we performed GSEA on DEGs from *LKB1* and *SMARCA4* tumours, using the same pipeline as the GSEA used for scRNA-seq data (**Table S4**). We then compared enriched biological processes between cell lines and tumours. Metabolic and cell cycle BP terms are common between *LKB1* and *SMARCA4* mutant lung tumours and cell line GSEA results, with genes that function in lipid metabolic process also enriched (**Figure 2B** and **Figure S2A**). We next compared the expression of enriched DEGs that comprise lipid metabolic process identified in GSEA of tumours between our scRNA-seq and cBioPortal bulk RNAseq datasets using heatmap plot (**Figure 2C,D** and **Table S5**). Most of the genes have expression patterns of LS-DEGs (**Figure 2C**), and display similar expression levels in *L*vsWT, *S*vsWT, and *LS*vsWT. There is individual contribution from both LKB1 and SMARCA4 to expression of genes linked to lipid metabolism but the majority of DEGs are impacted by both *LKB1* and *SMARCA4* loss. We also used our workflow to explore the LKB1-SMARCA4 transcriptional phenotype but used WT and *KRAS* mutant tumours as a control. When comparing *LKB1*, and *SMARCA4* mutant tumours to WT, *TP53*, and *KRAS* mutant tumours, we observed an inverse relationship with respect to correlation scores of enriched GO terms in *LKB1* and *SMARCA4* mutant tumours compared to WT, *KRAS* and *TP53* mutant tumours (**Table S6**). Furthermore, enriched GO terms in *LKB1* and *SMARCA4* mutant tumours exhibit similar enrichment scores in all three comparisons (**Table S6**). Therefore, these results suggest there is some transcriptional similarity between *LKB1* and *SMARCA4* mutant tumours (**Table S6**).

We next examined the expression profile of lipid metabolic process DEGs in *TP53*, *LKB1*, and *SMARCA4* early-stage and late-stage lung tumours (**Figure 2D**). Early stage *TP53*, *LKB1*, and *SMARCA4* tumours present with largely down-regulation of lipid metabolic process genes (**Figure 2D**). Interestingly, in late-stage tumours, we observed that *LKB1* and *SMARCA4* demonstrate strikingly similar expression profiles of genes involved in lipid metabolic process, specifically up-regulation of several genes (**Figure 2D**). Furthermore, this expression signature is different from late-stage *TP53* tumours and early-stage *LKB1* and *SMARCA4* tumours, suggesting that *LKB1* and *SMARCA4* mutant tumours exhibit similar phenotypes which manifest only in late-stage disease progression (**Figure 2D**). To determine if other pathways exhibit similar expression signature, we examined DEGs that comprised the cell cycle BP term, and we observed a similar gene expression pattern between *LKB1* and *SMARCA4* mutant tumours, with cell cycle genes largely up-regulated in late-stage tumours and in cell lines (**Figure S2B,C**). These results point to LKB1 and SMARCA4 functioning together to regulate tumourigenesis in lung cancer.

### SMARCA4 and LKB1 regulate gene expression in a linear pathway

The transcriptional similarity between our scRNA-seq data using cells in culture and bulk RNA-seq data from human lung tumours established a link between LKB1 and SMARCA4 in transcriptional regulation. We noticed that the majority of DEGs from scRNA-seq were classified as LS-DEGs, encompassing ∼50% of all genes detected in our scRNA analysis, indicating LKB1 similarly plays an important role in regulating gene expression as SMARCA4 (**Figure 1B**). Furthermore, L-DEGs outnumbered S-DEGs suggesting a greater requirement for LKB1 in regulating gene expression. Since LKB1 has been shown to bind and promote the ATPase activity of SMARCA4, and LKB1 localizes to both the nucleus and cytoplasm (15, 41, 42), we postulated that the transcriptional regulatory function of LKB1 could be mediated primarily through SMARCA4. To test this, we compared the expression LS-DEGs present in *L*vsWT to *S*vsWT using a scatter plot (**Figure 3A**). We noticed that LS-DEGs were similarly expressed in *L*vsWT compared to *S*vsWT, with a strong correlation of expression (R=0.69) (**Figure 3A**). Interestingly, our analysis revealed that only 454 genes out of 5528 (8.2% of genes) were inversely regulated, suggesting that LKB1 and SMARCA4 primarily co-regulate LS-DEGs in a linear pathway (**Table S1**).

**Figure 3.**
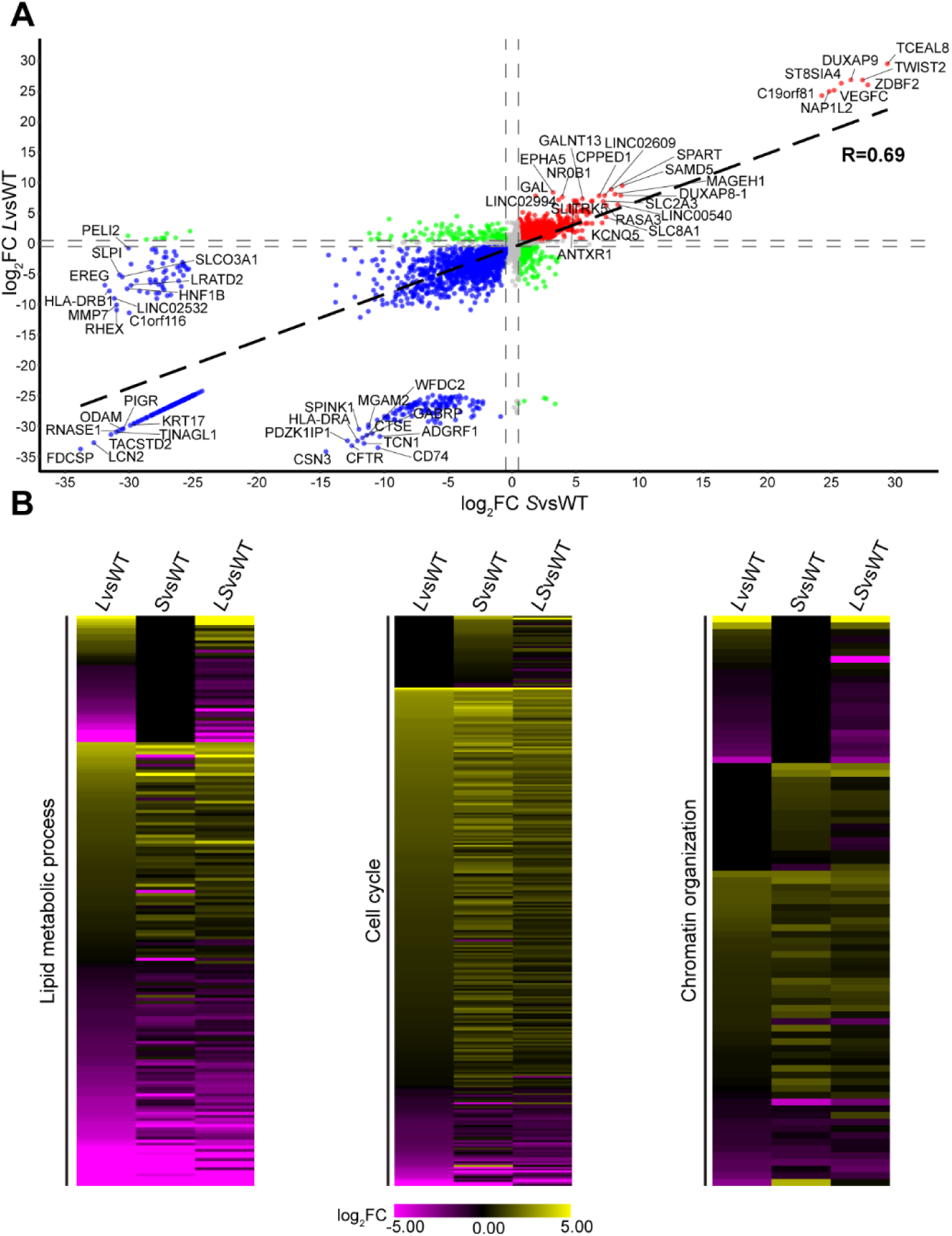
LKB1 and SMARCA4 regulate a similar expression program. **A**) Scatter plot of DEG expression in *L*vsWT compared with *S*vsWT. Red represents up-regulated genes above log_2_FC=0.5 threshold and blue represents down-regulated genes below log_2_FC=-0.5 threshold, while grey represents genes within log_2_FC 0.5 and −0.5 threshold. Green dots represent inversely related genes. Black dashed lines represent correlation trend line (R=0.69) and dashed grey lines represent log_2_FC 0.5 and −0.5 thresholds. Top and bottom 10 genes beyond the thresholds from *L*vsWT and *S*vsWT are labelled. **B**) Heatmap plot displaying expression of DEGs identified in lipid metabolic process, cell cycle, and chromatin organization from GSEA analysis of L-DEGs, S-DEGs, and LS-DEGs in *L*vsWT, *S*vsWT, and *LS*vsWT. Yellow represents high expression, and purple represents low expression with scale of −5 to 5 log_2_FC. DEGs not detected or have no change in expression are represented in black.

To evaluate the consequences that *LKB1* and *SMARCA4* loss have on gene expression, we visualized the expression pattern of DEGs identified in our GSEA for lipid metabolic process, cell cycle, and chromatin organization in *L*vsWT, *S*vsWT, and *LS*vsWT using heatmap plots (**Figure 3B** and **Table S7**). We observed that L-DEGs involved in lipid metabolic process are generally down-regulated and are similarly expressed in *L*vsWT compared to *LS*vsWT. LS-DEGs that function in lipid metabolic process displayed comparable differential expression when either *LKB1* or *SMARCA4* is mutated, with equivalent number of genes up and down-regulated, suggesting loss of either *LKB1* or *SMARCA4* produced similar transcriptional consequences on genes implicated in lipid metabolic process (**Figure 3B**). Likewise, *LKB1* or *SMARCA4* mutations had the same effects to expression of genes connected to cell cycle and chromatin organization (**Figure 3B**). Loss of either *LKB1* or *SMARCA4* caused up-regulation of cell cycle genes, and the magnitude of up-regulation was similar between *LKB1* loss and *SMARCA4* loss. We also noticed a similar profile with genes that function in chromatin organization (**Figure 3B**). These results highlight the phenotypic similarity loss of *SMARCA4* or *LKB1* have on gene expression related to diverse biological processes and that LKB1 and SMARCA4 function in a linear pathway.

The similar transcriptional profile observed in multiple biological processes between *LKB1* and *SMARCA4* loss suggested that LKB1 and SMARCA4 largely function together to regulate gene expression (19). To validate our results, we applied an unbiased approach where we examined the expression of genes within our scRNA-seq dataset that annotated to lipid metabolic process, cell cycle, and chromatin organization from the Gene Ontology Resource (43, 44). We reasoned that, if LKB1 and SMARCA4 function in a linear pathway with respect to gene expression regulation, most of the annotated genes from these biological processes would be characterized as LS-DEGs, and display similar expression profiles in *L*vsWT, *S*vsWT, and *LS*vsWT, like the DEGs from the GSEA analysis (**Figure 3B**). We visualized the expression profile of genes annotated to lipid metabolic process, cell cycle, and chromatin organization BP terms and observed that *LKB1* or *SMARCA4* loss results in similar transcriptional phenotype (**Fig. S3a** and **Table S7**). The majority of DEGs classified as LS-DEGs, and LKB1 and SMARCA4 provided distinct contributions to gene expression in these biological processes (**Figure S3A,B**). In lipid metabolic process, L-DEGs outnumber S-DEGs (n=78 vs n=45) suggesting a greater requirement for LKB1 in lipid metabolism. Similarly, there is a greater contribution from SMARCA4 in chromatin remodelling with S-DEGs outnumbering L-DEGs (n=55 vs n=40).

To further delve into the overlapping function of LKB1 and SMARCA4, we also examined the expression of genes annotated to immune response BP term (**Figure S1B,C**). LKB1 has been implicated in immune response as human *LKB1* mutant tumours are characterized to have low immunogenicity, and *LKB1* loss is associated with high inflammatory characteristics(45, 46).Tumours deficient of *LKB1* display gene expression profiles indicative of a suppressive immune tumour microenvironment, with observations of reduced infiltration of CD8+/CD4+ T-cells (12). This phenotype can be recapitulated with loss of AMPK activity, linking *LKB1* loss with diminished AMPK activity. This coupled with the observation that LS-DEGs showed strong enrichment for genes involved in immunity suggested that SMARCA4 and LKB1 collaborated to regulate expression of immune genes (**Figure 1D** and **Figure S1B,C**). We found L-DEGs enriched in immune function are down-regulated (**Figure S1B**) while S and LS-DEGs show similar number of up and down-regulated genes. Expression of most immune response genes are dependent on both *LKB1* and *SMARCA4* (n=548 out of n=985 DEGs, ∼55.6%), indicating LKB1 and SMARCA4 may collaborate to regulate expression of immune response genes.

### SMARCA4 regulates expression of LKB1-associated genes

Our results point to LKB1 and SMARCA4 regulating expression of genes involved in LKB1 pathways. If this is true, we predict that SMARCA4 would also regulate expression of genes known to be associated or be substrates of LKB1. Hence, we curated a list of LKB1-associated genes, specifically metabolic regulators, chromatin remodelling enzymes, epigenetic modifying enzymes, or genes differentially expressed in *LKB1* mutant context in multiple systems (mouse and human cell lines and tumours) (2, 28, 29, 47). We examined the expression of LKB1-associated genes in *L*vsWT, *S*vsWT and *LS*vsWT and interestingly, we observed that the majority of DEGs classified as LS-DEGs, indicating SMARCA4 plays an important role in regulating transcription of LKB1-associated genes (**Figure 4A** and **Table S8**). Furthermore, the expression of LKB1-associated genes was remarkably similar between *L*vsWT, LvsWT, and L*S*vsWT, suggesting that LKB1 and SMARCA4 work together to regulate expression of genes specifically implicated by LKB1 (**Table S8**). These results suggests that LKB1 and SMARCA4 may function in a linear pathway to regulate gene expression (**Figure 4B**).

**Figure 4.**
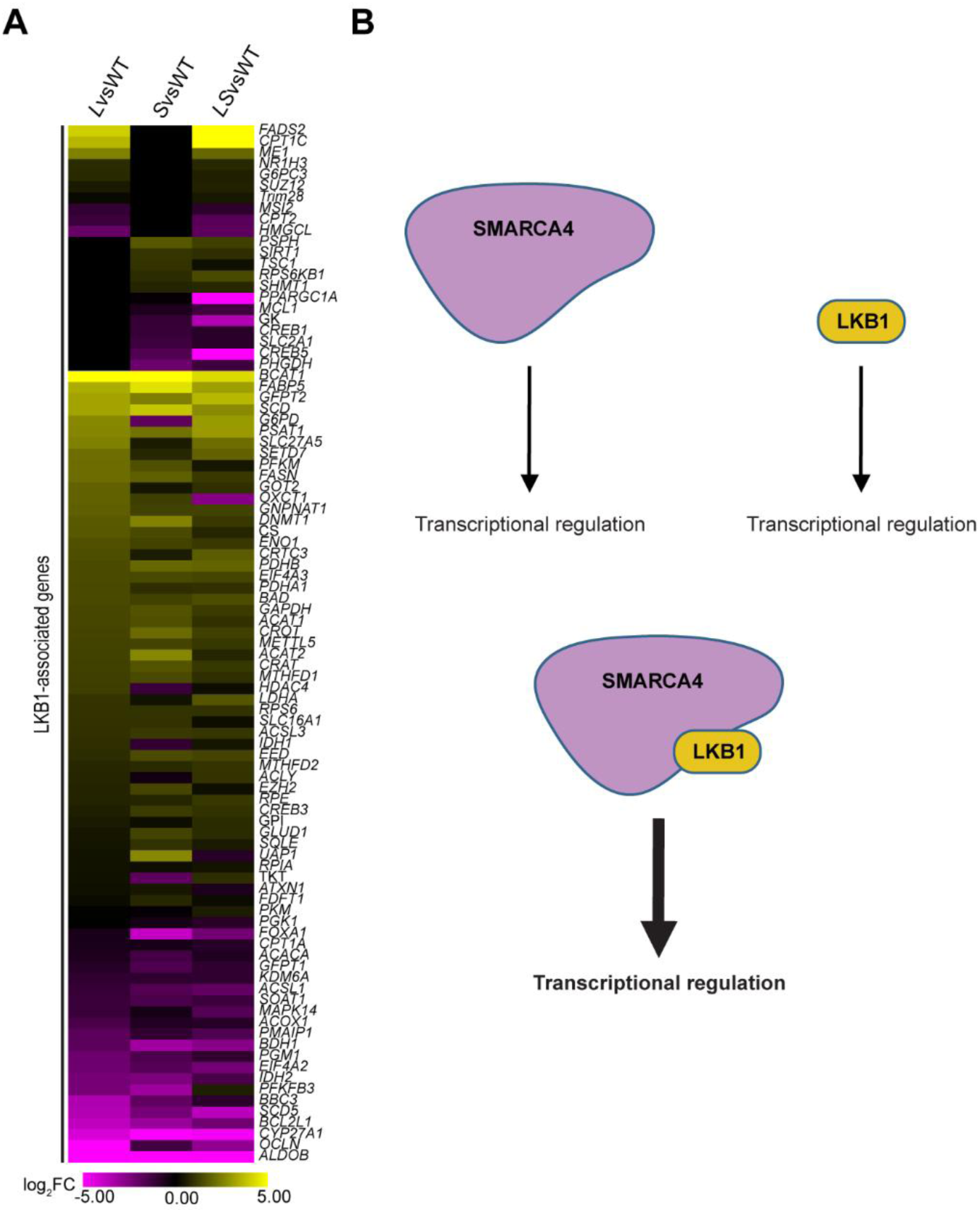
SMARCA4 regulates expression of LKB1-associated genes. **A**) Heatmap plot showing expression profile of LKB1-associated genes in LvsWT, SvsWT, and LSvsWT. Yellow represents high expression, and purple represents low expression with scale of −5 to 5 log2FC. DEGs not detected or have no change in expression are represented in black. b) SMARCA4 and LKB1 have independent and cooperative roles in transcriptional regulation.

### LKB1 functions with SMARCA4 to regulate lipid metabolism

During a state of energy imbalance, when AMP:ATP ratio is high, LKB1 activates AMPK, promoting increased lipid catabolism and reducing lipid anabolism. This phenotype is observed in *LKB1* mutant tumours, where *LKB1* catalytic activity is lost, tumours exhibit increased lipid accumulation, related to the diminished AMPK activity (48). To determine the impact SMARCA4 has on lipid metabolism, we focused on genes that comprised the lipid metabolic process BP term from GSEA analysis of *LKB1* and *SMARCA4* tumours and examined expression in scRNA-seq dataset (**Figure 2B**). Specifically, we investigated how SMARCA4 influences LKB1-mediated regulation of AMPK, which is crucial for controlling lipid metabolism (9, 10, 49), DEGs were then analysed using ExpressAnalyst (50) to further classify genes into anabolic (FA biosynthesis) and catabolic (FA β-oxidation, PPAR) processes (**Table S9**). We then made protein interaction network maps with NetworkAnalyst using the STRING database (51, 52) to visualize the pathway interactions (**Figure 5**). Genes involved in FA biosynthesis were upregulated when *LKB1* or *SMARCA4* are mutated, consistent with *LKB1* loss leading to upregulation of FA biosynthesis (53). We also observed SREBF1, the gene coding for SREBP1, is upregulated and transcriptionally regulated by LKB1 and SMARCA4 (Table 1). Conversely, genes involved in β-oxidation of FAs were down-regulated, with LKB1 and SMARCA4 contributing to regulation of β-oxidation genes (**Table S9** and **Figure 5**). We also observed peroxisome proliferator-activated receptors (PPARα and PPARγ), nuclear receptors that bind fatty acids and regulate gene expression involved in β-oxidation, are down-regulated (**Table S9** and **Figure 5**) (54). These results imply that LKB1 or SMARCA4 cooperate to regulate FA biosynthesis and β-oxidation, and that loss of *SMARCA4* displays a correlative gene expression pattern to *LKB1* loss.

**Figure 5.**
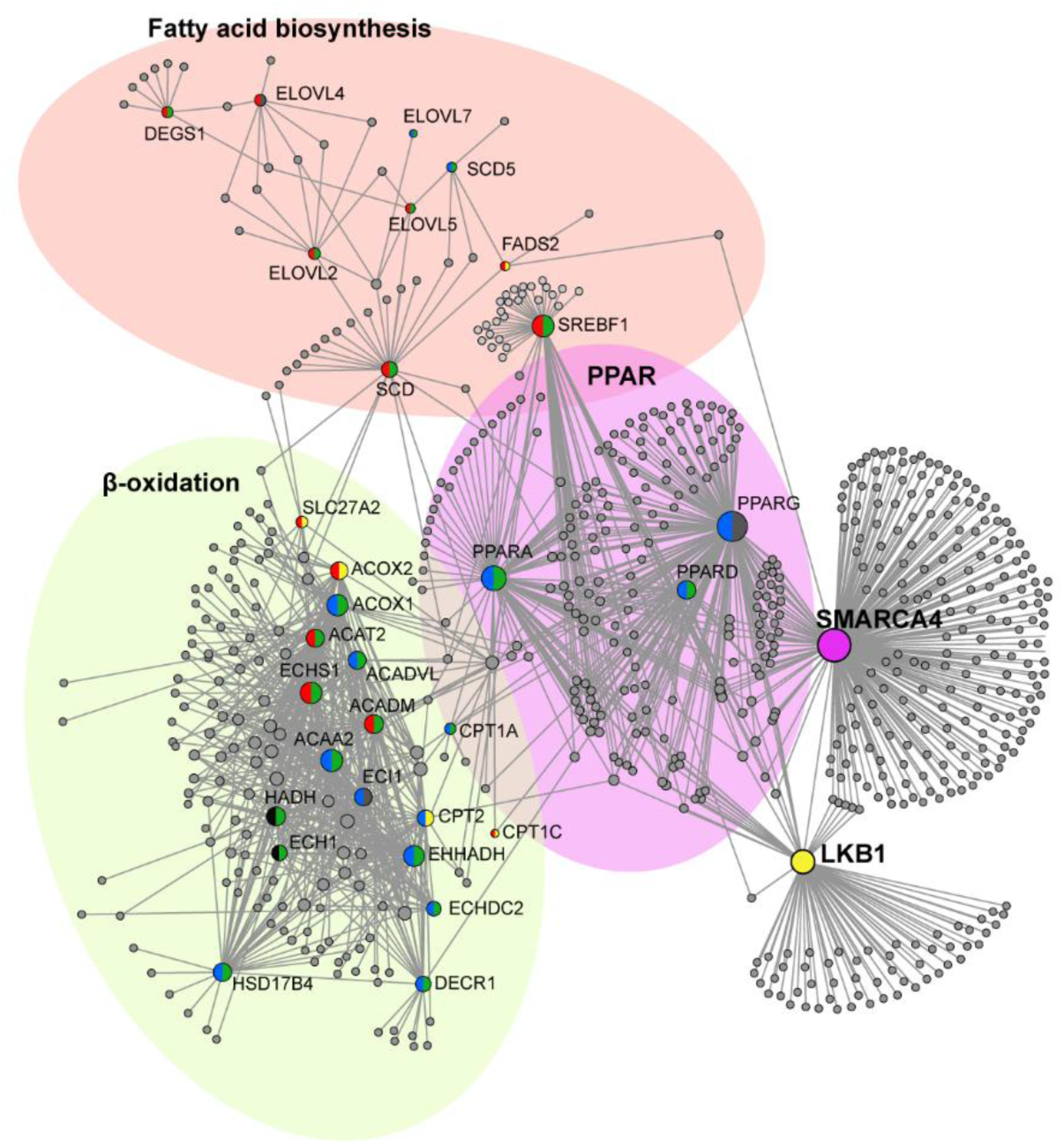
LKB1 and SMARCA4 collaborate to regulate lipid metabolism. Network maps of STRING interactions of genes involved in FA biosynthesis, PPAR, and β-oxidation. Nodes represent genes identified in GSEA. Node semi-circle colours represent up- or down-regulation and DEG classification as follows: yellow=L-DEGs, purple=S-DEGs, green=LS-DEGs, red=up-regulated, blue-down=regulated, black=in-consistent expression profile between *L*vsWT, *S*vsWT, and *LS*vsWT, and grey=classification other than L-DEG, S-DEG, or LS-DEG. Whole yellow circle represents LKB1, whole purple circle represents SMARCA4, and whole grey circles represent protein interactors experimentally determined. Generic PPI, with medium confidence stringency used (600) and experimental evidence (52).

## Discussion

Our results suggest that LKB1 and SMARCA4 tumour suppressor functions are interconnected, and that LKB1 and SMARCA4 may function together to regulate gene expression implicated in diverse biological processes in lung cancer. We found that loss of either *LKB1* or *SMARCA4* results in similar transcriptional profiles, with genes involved in metabolism, chromatin organization, cell cycle, and immune response showing similar gene expression profiles. Finally, we validated our scRNA-seq results using human lung tumours bulk RNA-seq data from the cBioPortal tumour database, implicating the LKB1-SMARCA4 interaction in lung tumourigenesis.

We propose a model for LKB1-SMARCA4 mediated transcriptional regulation based on our findings and suggest LKB1 has two master regulatory functions: the first is as a master regulatory kinase regulating energy metabolism and availability of cofactors used for chromatin remodelling, and the second as a transcriptional regulator in complex with SMARCA4 in the nucleus to regulate chromatin remodelling and gene expression (**Figure 6**) (19).

**Figure 6.**
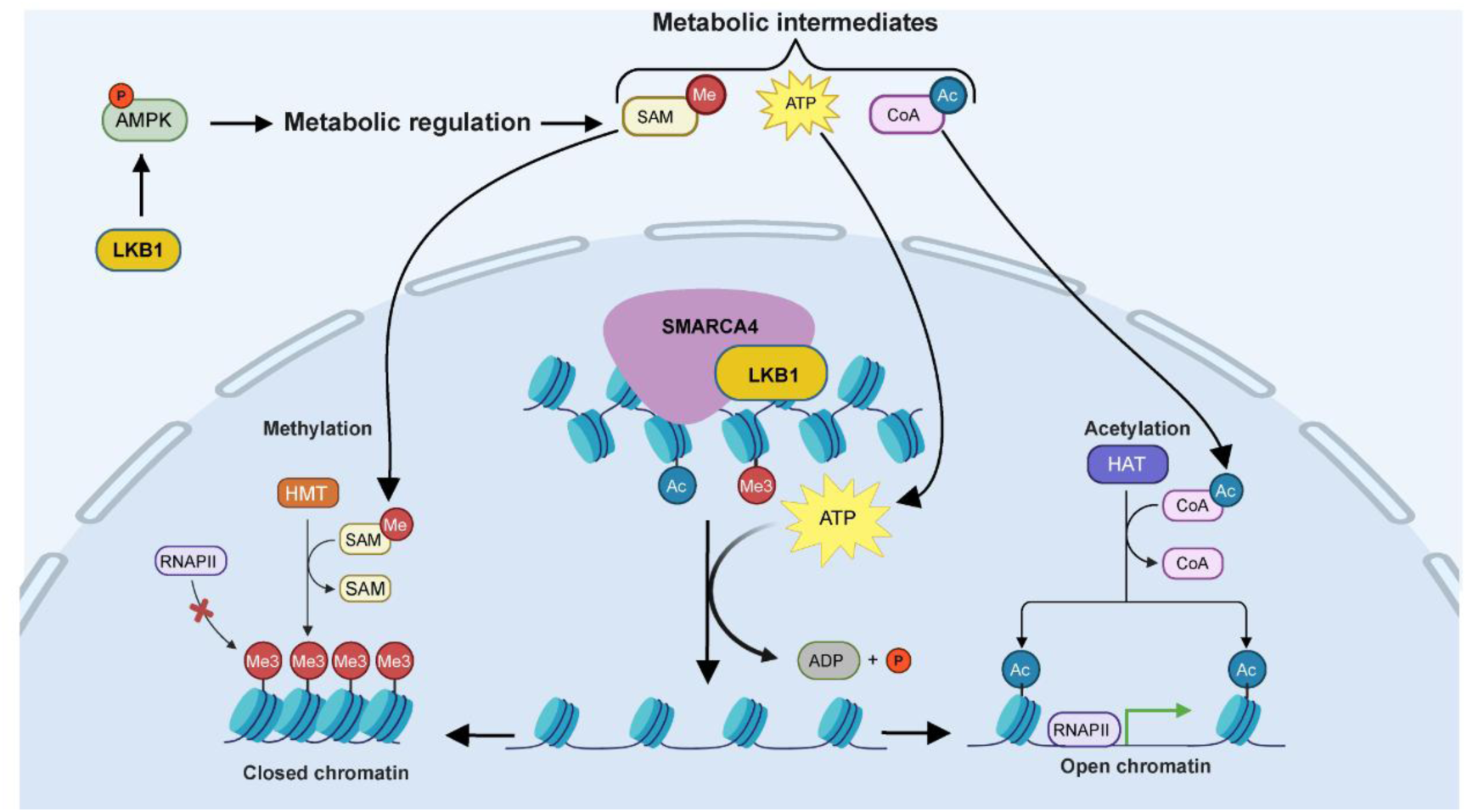
LKB1 and SMARCA4 cooperate to regulate chromatin remodeling and gene expression. Proposed model for LKB1 and SMARCA4 regulation of gene expression. LKB1 functions in the cytoplasm to regulate metabolic pathways through AMPK phosphorylation and activation. Metabolic intermediates ATP, acetyl-CoA, and SAM, translocate to the nucleus where LKB1 stimulates the ATPase activity of SMARCA4 at target loci determined by histone modifications to disrupt nucleosomes and modify chromatin organization. The remodeled chromatin can then be further modified through epigenetic modifications using metabolic intermediates from LKB1 regulated pathways as cofactors. Histone methyltransferases (HMTs) can use SAM to promote histone methylation (ME3), creating closed chromatin and blocking transcription. Histone acetyltransferases (HATs) can use acetyl-CoA to acetylate histones (Ac), creating open chromatin and promote transcription. Image created using BioRender and processed using adobe illustrator.

*LKB1* lung tumours exhibit increased lipid accumulation due to attenuation of AMPK signalling. In ATP deprived state (when AMP:ATP ratio is high), LKB1 phosphorylates and activates AMPK leading to inhibition of anabolic pathways, and promotion of catabolic pathways (8). When *LKB1* is mutated, genes involved in lipid biosynthesis are up-regulated, and genes involved in β-oxidation are down-regulated (**Figure 5**). FA synthesis is regulated by in part by AMPK phosphorylation of SREBP1, which is a transcription factor regulating expression of genes involved in FA synthesis (20, 21). We observe genes involved in FA biosynthesis are also up regulated when *SMARCA4* is lost, even in the presence of LKB1, suggesting that SMARCA4 is epistatic to LKB1 in regulating gene expression of FA biosynthesis genes (**Figure 5**). SMARCA4 has also been shown to play a role in expression of genes involved in FA biosynthesis (**Figure 5**) (55). In hepatocytes, SREBP1 was shown to bind SMARCA4-dependent SWI/SNF complex suggesting FA biosynthesis gene expression is regulated by LKB1 and SMARCA4 in a linear pathway (55).

LKB1 has been shown to interact with SMARCA4 in the nucleus (15) and loss of this interaction leads to global downregulation of gene expression (19). Furthermore, our lab has shown that LKB1 binds and promotes the ATPase-dependent chromatin remodelling function of SMARCA4 (15). Therefore, we speculate that the striking correlated gene expression profiles that results when either *LKB1* or *SMARCA4* is mutated may be due to the absence of the LKB1-SMARCA4 physical interaction (15). Individual functions of LKB1 and SMARCA4 in regulating gene expression are observed from L-DEGs and S-DEGs implicated in diverse pathways, but we consistently view LS-DEGs are the predominant classification (**Figure 4B** and **Figure S3A**). Our findings support the hypothesis that the primary mechanism by which LKB1 regulates gene expression is through the previously observed physical interaction with SMARCA4; whereby the LKB1-SMARCA4 interaction regulates the recruitment of the transcriptional and epigenetic regulators to gene regulatory elements mediated by binding with SMARCA4, promoting the ATPase activity and chromatin remodelling function of SMARCA4 (15).

LKB1 metabolic regulation can also impact the chromatin landscape independently of SMARCA4 (17, 56–58). Therefore, it is possible that our observations could be through independent but collaborative pathways integrating at epigenetic modification pathways, in addition to the previously observed LKB1-SMARCA4 interaction(17, 56–58). Metabolic pathways produce intermediates such as ATP, acetyl-CoA, and SAM among others, that are used as cofactors for chromatin remodelling enzymes that regulate global changes in chromatin organization and consequently gene expression (1, 2, 59). The metabolic state of cells influences the abundance of these cofactors. For example, lipid metabolism drastically impacts levels of ATP and acetyl-CoA, through regulation of FA biosynthesis and β-oxidation (3). Conversely, SAM production is dependent on the serine-glycine one carbon cycle (2). *LKB1* loss is associated with global chromatin remodelling, and increased histone methylation through up-regulation of SAM production (2, 28).It is possible that metabolic intermediates from LKB1 regulated pathways are utilized by chromatin remodelling complexes to regulate epigenetic modifications. Moreover, chromatin remodelling complexes can interact with epigenetic enzymes, regulating histone modifications, and in this way, LKB1 can regulate epigenetic modifications indirectly from SMARCA4 (60, 61). We suggest that LKB1 acts as a master regulator of gene expression through metabolic pathway regulation, production of epigenetic cofactors, and as a regulator of chromatin remodelling via the physical interaction with SMARCA4. Since LKB1 can traverse the nuclear membrane (41) where it can then be recruited to transcriptional machinery through direct binding within the nucleus, LKB1 could function as a integrator of the metabolic state of cells to responses in gene expression, impacting production of ATP and other metabolic intermediates utilized by SMARCA4 and the chromatin remodelling machinery to regulate chromatin structure and consequently gene expression (**Figure 5**).

In summary, we show that *LKB1* and *SMARCA4* mutants exhibit similar expression profiles in lung cancer cell lines and in human lung tumours and this interaction contributes to the regulation of diverse pathways in lung cancer. We propose a model where the tumour suppressors LKB1 and SMARCA4 function together to prevent lung cancer through both direct and indirect mechanisms converging to regulate metabolism and gene expression. Our results provide evidence that implicates LKB1 as a master regulator, serving as an interface between metabolic pathways and gene expression. We propose this function of LKB1 is mediated in part through physical interaction with SMARCA4, positioning LKB1 as a nexus between metabolism and gene expression.

## Methods

### Workflow to identify LKB1 (L) SMARCA4 (S) and LKB1-SMARCA4 (LS) DEGs

Single-cell RNA-seq data (scRNA-seq) mined and process as in our previous study (32). All DEGs detected from scRNA-seq were identified and log_2_FC expression level for each gene was determined with Calu-3 cells serving as a control: H460 vs. Calu-3 (*L*), H1299 vs. Calu-3 (*S*), and A549 vs. Calu-3 (*LS*). Classification of DEGs was as follows: L-DEGs: detected in *L* and *LS,* not in *S*. S-DEGs: detected in *S* and *LS,* not in *L*. LS-DEGs: detected in *L*, *S*, and *LS*. Each classification represented a gene set for gene set enrichment analysis.

### Data processing and gene set enrichment analysis

For gene set enrichment analysis, we used <u>g:Profiler</u> (62) to identify enriched Gene Ontology (GO) terms at the category of biological process (BP) using default settings. Results from up-, down-regulated, and complete gene sets were combined, and redundant BP terms were filtered for highest intersection size. Enriched BP terms were then compared to each other using a similarity matrix based on overlapping gene IDs. Hierarchical clustering was performed using enriched BP terms using hierarchical k-means clustering (silhouette method) to identify unique GeneIDs for each cluster. The similarity matrices and hierarchical clustering were generated using RStudio. Unique gene IDs for each cluster were then analyzed using <u>ExpressAnalyst</u> (63)network enrichment tool using Panther database to generate enriched BP terms for dot plots, with only terms >3 Gene IDs (hits) chosen and only statistically significant term results collected (FDR-corrected p-value<0.05). Genes that annotated to lipid metabolic process, chromatin organization, cell cycle, and immune response GO terms were obtained from the Gene Ontology Resource (43, 44).

### Plots

Venn diagrams were generated using Ghent University <u>Venn diagram generator web tool</u>. Heatmap plots were generated using <u>Broad Institute Morpheus web tool</u>. Volcano and dot plots were generated using ggplot2 in RStudio. Network maps were made from <u>NetworkAnalyst</u> (51) and processed using Adobe Illustrator.

### Correlation analysis using scRNA-seq and bulk RNA-seq lung cancer datasets

We obtained a Transcripts Per Million (TPM)-normalized mRNA expression matrix of 517 bulk RNA-seq datasets from lung adenocarcinoma (LUAD) patients identified in The Cancer Genome Atlas (TCGA) and Broad Genome Data Analysis Center (GDAC) Firehose in the cBioPortal for Cancer Genomics database (37). For the correlation analysis, we selected 66 datasets based on a mutation status of three tumour suppressor genes, *LKB1, SMARCA4, and TP53*, and divided into four subgroups that are represented genotypes similar to the cell lines used for scRNA-seq; Calu-3 represented by *TP53* tumours (51), H460 represented by *LKB1* tumours (12), H1299 represented by *SMARCA4* tumours (2), and A549 represented by *LKB1,SMARCA4* tumours (1). A TPM-normalized mRNA expression matrix of the 66 subgroup datasets was used for hierarchical clustering analysis following a principal component analysis for dimension reduction. Finally, gene expression-based relationships among the four lung cancer patient subgroups (bulk RNA-seq) as well as the four lung cancer cell lines (scRNA-seq) were respectively identified using the Pearson-correlation method. In addition, we prepared two separate TPM-normalized mRNA expression matrixes from bulk RNA-seq datasets of the LUAD patients for 299 early-stage samples (237 wild type (WT), 7 *LKB1*-mutated, 3 *SMARCA4*-mutated, and 51 *TP53*-mutated) and 83 late-stage samples (61 WT, 2 *LKB1*-mutated, 2 *SMARCA4*-mutated, and 18 *TP53*-mutated).

A DEA was performed on each of the two mRNA expression matrixes to detect marker genes significantly up or down regulated in mutated samples of each tumour suppressor gene at the two different lung cancer stages when compared to WT samples. To select significant marker genes, we applied identical parameters that were used for the DEA of scRNA-seq datasets.

All the figures were processed and assembled using Adobe Illustrator.

## Acknowledgements

The authors would like to thank members of the Marignani lab for their support. This work was supported by the Cancer Research Society, The Canadian Cancer Society Diane Campbell designated research fund (grant #706202). The authors acknowledge that *Dalhousie University sits on Mi’kma’ki, the ancestral and unceded territory of the Mi’kmaq People. We are all Treaty People. We recognize that African Nova Scotians are a distinct people whose histories, legacies and contributions have enriched that part of Mi’kma’ki known as Nova Scotia for over 400 years*.

## Data availability

Data analysed in this study was obtained from the NCBI Gene Expression Omnibus database (GEO ID: GSE183590). Human lung tumour data was obtained from TCGA and GDAC Firehose dataset from cBioPortal (38).

## Conflict of interest statement

The authors declare no conflicts of interest.

---

**Fig. S1.**
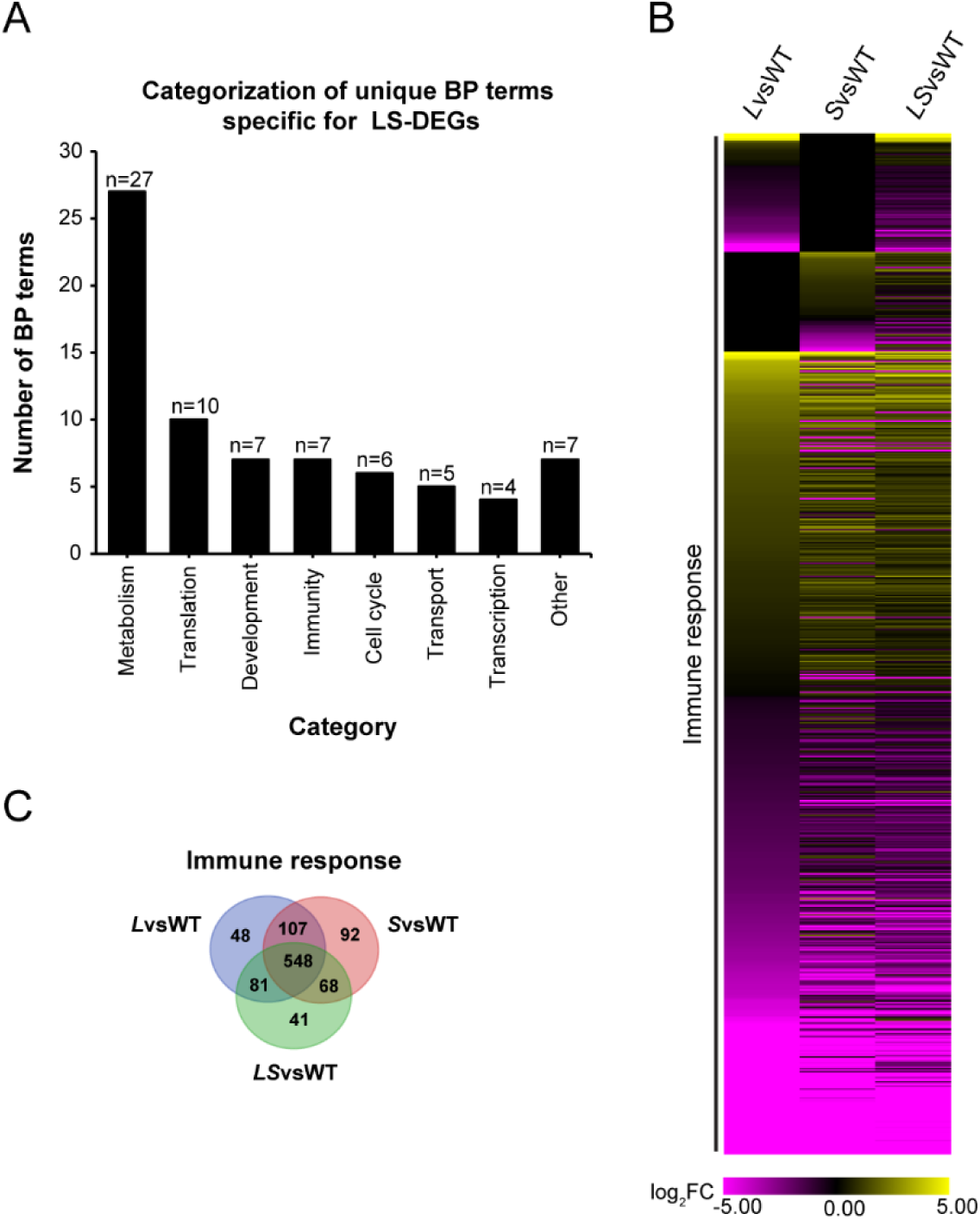
LKB1 and SMARCA4 function together to regulate expression of immune response genes. **A**) Bar plot showing the categorization of BP terms for GSEA analysis of LS-DEGs. **B**) Heatmap plot representing expression of genes annotated to immune response BP term from the GO consortium in *L*vsWT, *S*vsWT, and *LS*vsWT. Yellow represents high expression, and purple represents low expression with scale of −5 to 5 log_2_FC. DEGs not detected or have no change in expression are represented in black. **C**) Venn diagram depicting classification of genes that comprise immune response BP from scRNA-seq data.

**Fig. S2.**
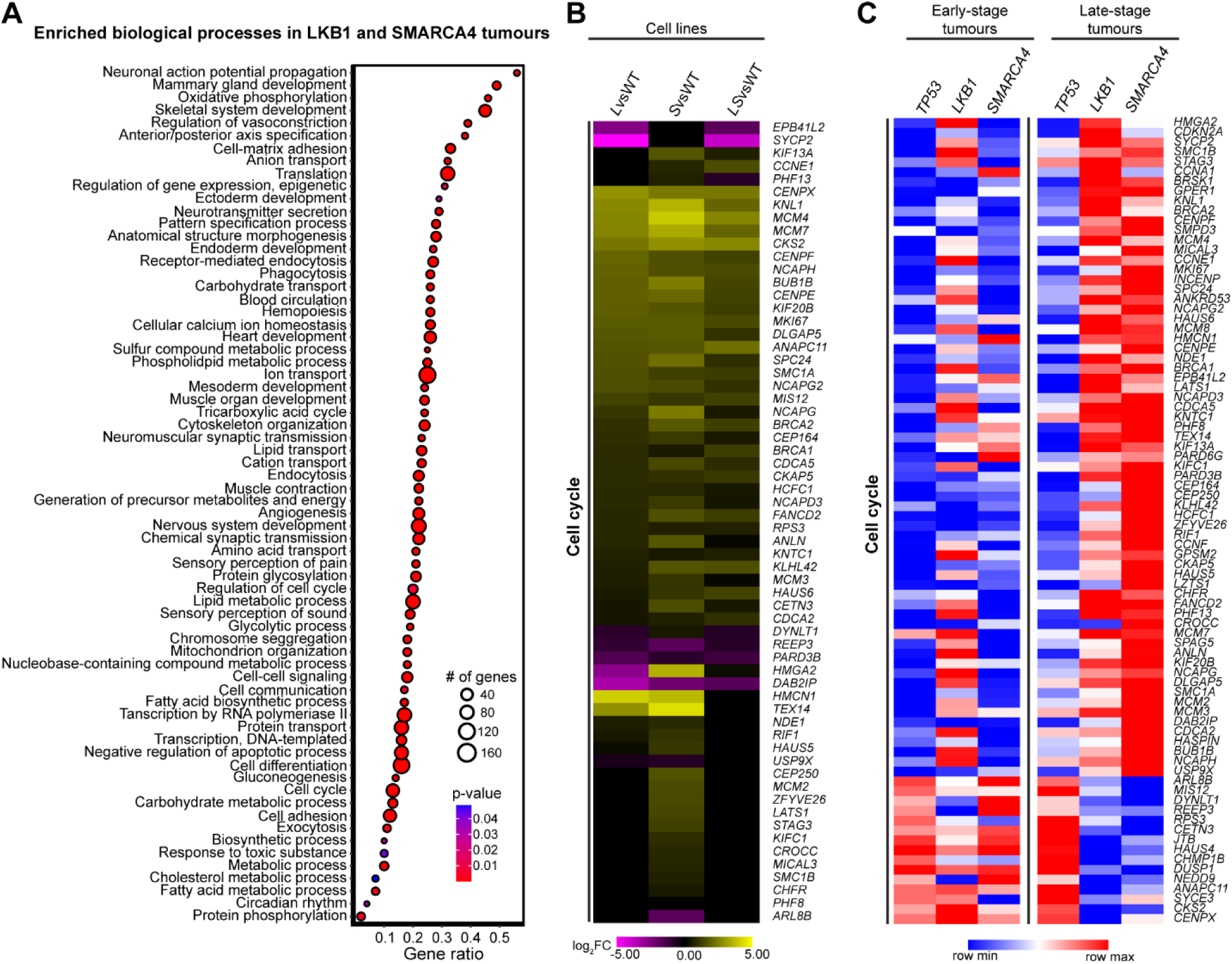
LKB1 and SMARCA4 regulate similar biological processes in human lung tumours. **A)** Dot plot of enriched BP terms from GSEA analysis of *LKB1* and *SMARCA4* tumour DEGs. Gene ratio represents number of genes/term size. Dot size represents the number of genes identified in each BP term, colour represents p-value with red representing low p-value and blue represents high p-value. Only BP terms that are statistically significant (*P*<0.05) and have >3 genes are shown. **b,c**) Heatmap plots showing expression of DEGs identified in GSEA analysis from *LKB1* and *SMARCA4* tumour DEGs involved in cell cycle process. **B**) The heatmap plot represents expression of cell cycle genes from *L*vsWT, *S*vsWT, and *LS*vsWT using scRNA-seq data (purple/yellow). Yellow represents high expression, and purple represents low expression with scale of −5 to 5 log_2_FC. DEGs not detected or have no change in expression are represented in black. **C**) The heatmap plot represents expression of cell cycle genes from *TP53*, *LKB1*, and *SMARCA4* early-stage and late-stage tumours (red/blue). Red represents high expression, and blue represents low expression with scale of row min row max.

**Fig. S3.**
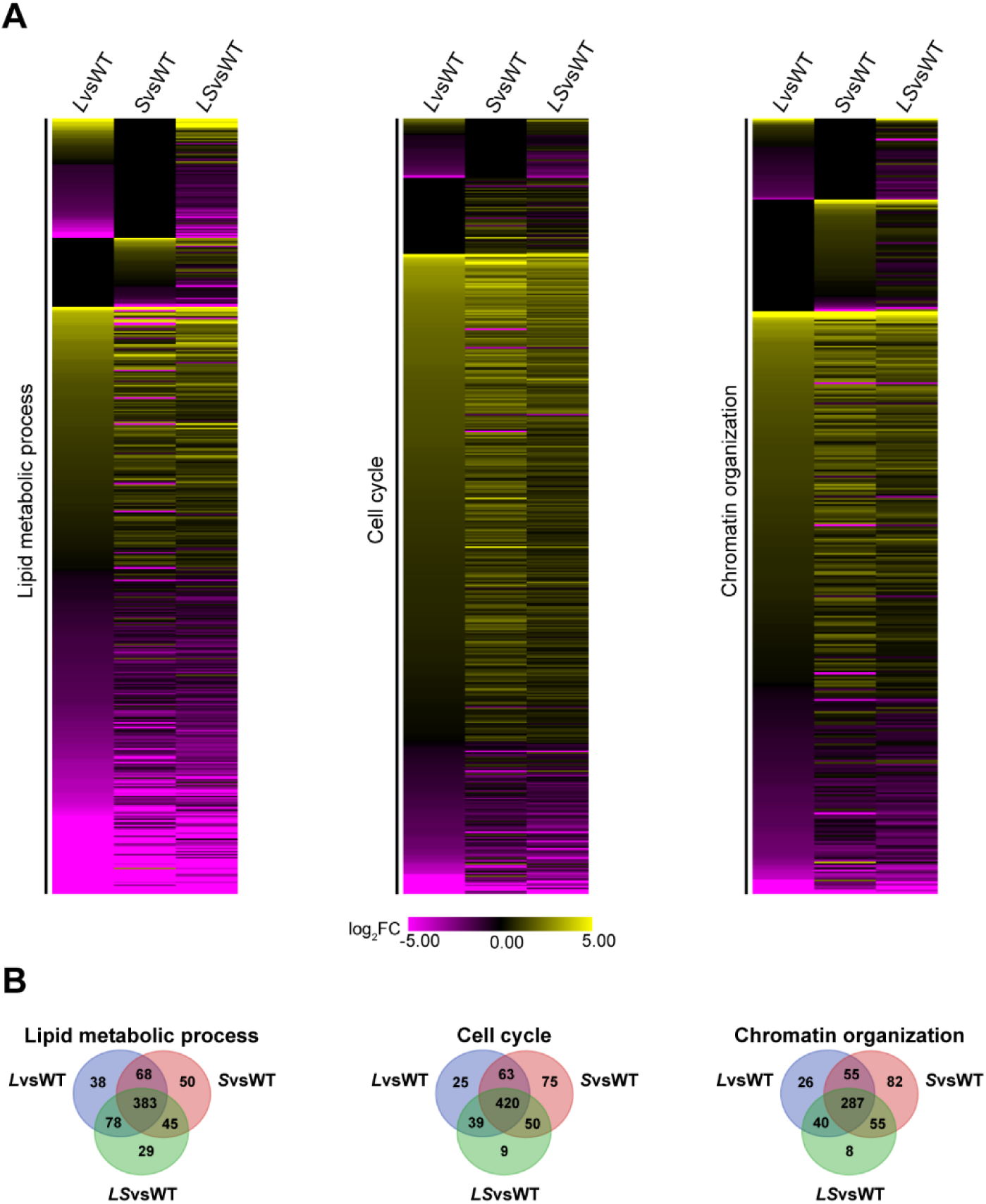
LKB1 and SMARCA4 collaborate to regulate expression of genes in multiple pathways. **A**) Heatmap plot displaying expression of genes annotated to BP terms lipid metabolic process, cell cycle, and chromatin organization from the GO consortium in *L*vsWT, *S*vsWT, and *LS*vsWT. Only L-DEGs, S-DEGs, and LS-DEGs are shown. Yellow represents high expression, and purple represents low expression with scale of −5 to 5 log_2_FC. DEGs not detected or have no change in expression are represented in black. **B**) Venn diagrams showing classification of DEGs from BP terms in **A** (lipid metabolic process, chromatin organization, and cell cycle).

## Supplementary Information

**Table S1.** Log_2_FC expression of DEGs from scRNA-seq dataset in *L*vsWT, *S*vsWT, and *LS*vsWT

**Table S2.** gProfiler and ExpressAnalyst results L-DEGs, S-DEGs, LS-DEGs

**Table S3.** cBioPortal tumor mutation information, Log_2_FC expression of LKB1 SMARCA4 tumour DEGs

**TableS4.** gProfiler and ExpressAnalyst results for LKB1 and SMARCA4 tumour DEGs

**Table S5.** Heatmap plots for human lung tumor dataset

**Table S6.** TCGA Enriched BP Terms in LKB1 and SMARCA4 mutant tumours

**Table S7.** Heatmap plots scRNA-seq dataset

**Table S8.** Log_2_FC expression of LKB1-associated genes in LvsWT, SvsWT, and LSvsWT

**Table S9.** Genes involved in FA biosynthesis, PPAR, and β-oxidation

## Notes

### Competing Interest Statement

The authors have declared no competing interest.

## References

1. Gut P, Verdin E. The nexus of chromatin regulation and intermediary metabolism. Nature. 2013;502(7472):489–98.

2. Kottakis F, Nicolay BN, Roumane A, Karnik R, Gu H, Nagle JM, et al. LKB1 loss links serine metabolism to DNA methylation and tumorigenesis. Nature. 2016;539(7629):390–5.

3. Sutendra G, Kinnaird A, Dromparis P, Paulin R, Stenson TH, Haromy A, et al. A nuclear pyruvate dehydrogenase complex is important for the generation of acetyl-CoA and histone acetylation. Cell. 2014;158(1):84–97.

4. Bultman S, Gebuhr T, Yee D, La Mantia C, Nicholson J, GIlliam A, et al. A Brg1 null mutation in the mouse reveals functional differences among mammalian SWI/SNF complexes. Mol Cell. 2000;6:1287–95.

5. Kwon H, Imbalzano AN, Khavari PA, Kingston RE, Green MR. Nucleosome disruption and enhancement of activator binding by a human SW1/SNF complex [see comments]. Nature. 1994;370(6489):477–81.

6. Rodriguez-Nieto S, Sanchez-Cespedes M. BRG1 and LKB1: tales of two tumor suppressor genes on chromosome 19p and lung cancer. Carcinogenesis. 2009;30(4):547–54.

7. Marignani PA. LKB1, the multitasking tumour suppressor kinase. J Clin Pathol. 2005;58(1):15–9.

8. Bourouh M, Marignani PA. The Tumor Suppressor Kinase LKB1: Metabolic Nexus. Frontiers in Cell and Developmental Biology. 2022;10.

9. Hawley S, Boudeau J, Reid J, Mustard K, Udd L, Makela T, et al. Complexes between the LKB1 tumor suppressor, STRAD alpha/beta and MO25 alpha/beta are upstream kinases in the AMP-activated protein kinase cascade. J Biol. 2003;2:28.

10. Shaw R, Kosmatka M, Bardeesy N, Hurley R, Witters L, DePinho R, et al. The tumor suppressor LKB1 kinase directly activates AMP-activated kinase and regulates apoptosis in response to energy stress. Proc Natl Acad Sci USA. 2004;101:3329–35.

11. La Fleur L, Falk-Sörqvist E, Smeds P, Berglund A, Sundström M, Mattsson JS, et al. Mutation patterns in a population-based non-small cell lung cancer cohort and prognostic impact of concomitant mutations in KRAS and TP53 or STK11. Lung Cancer. 2019;130:50–8.

12. Gao Y, Päivinen P, Tripathi S, Domènech-Moreno E, Wong IPL, Vaahtomeri K, et al. Inactivation of AMPK Leads to Attenuation of Antigen Presentation and Immune Evasion in Lung Adenocarcinoma. Clin Cancer Res. 2022;28(1):227–37.

13. Medina P, Romero O, Kohno T, Montuenga L, Pio R, Yokota J, et al. Frequent BRG1/SMARCA4-inactivating mutations in human lung cancer cell lines. Human Mutation. 2008;29(5):617–22.

14. Schoenfeld AJ, Bandlamudi C, Lavery JA, Montecalvo J, Namakydoust A, Rizvi H, et al. The Genomic Landscape of SMARCA4 Alterations and Associations with Outcomes in Patients with Lung Cancer. Clin Cancer Res. 2020;26(21):5701–8.

15. Marignani PA, Kanai F, Carpenter CL. LKB1 associates with Brg1 and is necessary for Brg1-induced growth arrest. J Biol Chem. 2001;276(35):32415–8.

16. Hemminki A, Markie D, Tomlinson I, Avizienyte E, Roth S, Loukola A, et al. A serine/threonine kinase gene defective in Peutz-Jeghers syndrome. Nature. 1998;391(6663):184–7.

17. Scott KD, Nath-Sain S, Agnew MD, Marignani PA. LKB1 catalytically deficient mutants enhance cyclin D1 expression. Cancer Res. 2007;67(12):5622–7.

18. Kang H, Cui K, Zhao K. BRG1 Controls the Activity of the Retinoblastoma Protein via Regulation of p21CIP1/WAF1/SDI. Mol Cell Biol. 2004;24(3):1188–99.

19. Mével-Aliset M, Radu AG, Allard J, Blanchet S, Montellier E, Hainaut P, et al. Transcriptional regulation by LKB1 in lung adenocarcinomas: Exploring oxidative stress, neuroglial and amino acid signatures. Biochem Biophys Res Commun. 2025;755:151571.

20. Seo MS, Kim JH, Kim HJ, Chang KC, Park SW. Honokiol activates the LKB1-AMPK signaling pathway and attenuates the lipid accumulation in hepatocytes. Toxicol Appl Pharmacol. 2015;284(2):113–24.

21. Li Y, Xu S, Mihaylova MM, Zheng B, Hou X, Jiang B, et al. AMPK phosphorylates and inhibits SREBP activity to attenuate hepatic steatosis and atherosclerosis in diet-induced insulin-resistant mice. Cell Metab. 2011;13(4):376–88.

22. Li X, Egervari G, Wang Y, Berger SL, Lu Z. Regulation of chromatin and gene expression by metabolic enzymes and metabolites. Nature Reviews Molecular Cell Biology. 2018.

23. Juszczak F, Caron N, Mathew AV, Declèves AE. Critical Role for AMPK in Metabolic Disease-Induced Chronic Kidney Disease. Int J Mol Sci. 2020;21(21).

24. Lee WJ, Kim M, Park HS, Kim HS, Jeon MJ, Oh KS, et al. AMPK activation increases fatty acid oxidation in skeletal muscle by activating PPARalpha and PGC-1. Biochem Biophys Res Commun. 2006;340(1):291–5.

25. Salma N, Xiao H, Mueller E, Imbalzano AN. Temporal recruitment of transcription factors and SWI/SNF chromatin-remodeling enzymes during adipogenic induction of the peroxisome proliferator-activated receptor gamma nuclear hormone receptor. Mol Cell Biol. 2004;24(11):4651–63.

26. Nath-Sain S, Marignani PA. LKB1 catalytic activity contributes to estrogen receptor alpha signaling. Mol Biol Cell. 2009;20(11):2785–95.

27. DiRenzo J, Shang Y, Phelan M, Sif S, Myers M, Kingston R, et al. BRG-1 is recruited to estrogen-responsive promoters and cooperates with factors involved in histone acetylation [In Process Citation]. Mol Cell Biol. 2000;20(20):7541–9.

28. Pierce SE, Granja JM, Corces MR, Brady JJ, Tsai MK, Pierce AB, et al. LKB1 inactivation modulates chromatin accessibility to drive metastatic progression. Nat Cell Biol. 2021;23(8):915–24.

29. Zhang H, Fillmore Brainson C, Koyama S, Redig AJ, Chen T, Li S, et al. Lkb1 inactivation drives lung cancer lineage switching governed by Polycomb Repressive Complex 2. Nat Commun. 2017;8:14922.

30. Haberman N, Cheung R, Pizza G, Cvetesic N, Nagy D, Maude H, et al. Liver kinase B1 (LKB1) regulates the epigenetic landscape of mouse pancreatic beta cells. Faseb j. 2024;38(16):e23885.

31. Carretero J, Medina PP, Pio R, Montuenga LM, Sanchez-Cespedes M. Novel and natural knockout lung cancer cell lines for the LKB1/STK11 tumor suppressor gene. Oncogene. 2004;23(22):4037–40.

32. Kim J, Xu Z, Marignani PA. Single-cell RNA sequencing for the identification of early-stage lung cancer biomarkers from circulating blood. NPJ Genom Med. 2021;6(1):87.

33. Blanco R, Iwakawa R, Tang M, Kohno T, Angulo B, Pio R, et al. A gene-alteration profile of human lung cancer cell lines. Human Mutation. 2009;30(8):1199–206.

34. Medina PP, Carretero J, Ballestar E, Angulo B, Lopez-Rios F, Esteller M, et al. Transcriptional targets of the chromatin-remodelling factor SMARCA4/BRG1 in lung cancer cells. Hum Mol Genet. 2005;14(7):973–82.

35. Hendricks KB, Shanahan F, Lees E. Role for BRG1 in Cell Cycle Control and Tumor Suppression. Mol Cell Biol. 2004;24(1):362–76.

36. Gupta M, Concepcion CP, Fahey CG, Keshishian H, Bhutkar A, Brainson CF, et al. BRG1 Loss Predisposes Lung Cancers to Replicative Stress and ATR Dependency. Cancer Res. 2020;80(18):3841–54.

37. Cerami E, Gao J, Dogrusoz U, Gross BE, Sumer SO, Aksoy BA, et al. The cBio cancer genomics portal: an open platform for exploring multidimensional cancer genomics data. Cancer discovery. 2012;2(5):401–4.

38. de Bruijn I, Kundra R, Mastrogiacomo B, Tran TN, Sikina L, Mazor T, et al. Analysis and Visualization of Longitudinal Genomic and Clinical Data from the AACR Project GENIE Biopharma Collaborative in cBioPortal. Cancer Res. 2023;83(23):3861–7.

39. Gao J, Aksoy BA, Dogrusoz U, Dresdner G, Gross B, Sumer SO, et al. Integrative analysis of complex cancer genomics and clinical profiles using the cBioPortal. Sci Signal. 2013;6(269):pl1.

40. Skoulidis F, Byers LA, Diao L, Papadimitrakopoulou VA, Tong P, Izzo J, et al. Co-occurring genomic alterations define major subsets of KRAS-mutant lung adenocarcinoma with distinct biology, immune profiles, and therapeutic vulnerabilities. Cancer discovery. 2015;5(8):860–77.

41. Nezu J, Oku A, Shimane M. Loss of cytoplasmic retention ability of mutant LKB1 found in Peutz-Jeghers syndrome patients. Biochem Biophys Res Commun. 1999;261(3):750–5.

42. Boudeau J, Baas AF, Deak M, Morrice NA, Kieloch A, Schutkowski M, et al. MO25alpha/beta interact with STRADalpha/beta enhancing their ability to bind, activate and localize LKB1 in the cytoplasm. Embo J. 2003;22(19):5102–14.

43. Ashburner M, Ball C, Blake J, Botstein D, Butler H, Cherry J, et al. Gene ontology: tool for the unification of biology. The Gene Ontology Consortium. Nat Genet. 2000;25(1):25–9.

44. Aleksander SA, Balhoff J, Carbon S, Cherry JM, Drabkin HJ, Ebert D, et al. The Gene Ontology knowledgebase in 2023. Genetics. 2023;224(1).

45. Koyama S, Akbay EA, Li YY, Aref AR, Skoulidis F, Herter-Sprie GS, et al. STK11/LKB1 Deficiency Promotes Neutrophil Recruitment and Proinflammatory Cytokine Production to Suppress T-cell Activity in the Lung Tumor Microenvironment. Cancer Research. 2016.

46. Skoulidis F, Goldberg ME, Greenawalt DM, Hellmann MD, Awad MM, Gainor JF, et al. STK11/LKB1 Mutations and PD-1 Inhibitor Resistance in KRAS-Mutant Lung Adenocarcinoma. Cancer discovery. 2018;8(7):822–35.

47. Timilshina M, You Z, Lacher SM, Acharya S, Jiang L, Kang Y, et al. Activation of Mevalonate Pathway via LKB1 Is Essential for Stability of T(reg) Cells. Cell Rep. 2019;27(10):2948–61.e7.

48. Bhatt V, Khayati K, Hu ZS, Lee A, Kamran W, Su X, et al. Autophagy modulates lipid metabolism to maintain metabolic flexibility for Lkb1-deficient Kras-driven lung tumorigenesis. Genes Dev. 2019;33(3-4):150–65.

49. Lizcano JM, Goransson O, Toth R, Deak M, Morrice NA, Boudeau J, et al. LKB1 is a master kinase that activates 13 kinases of the AMPK subfamily, including MARK/PAR-1. EMBO J. 2004;23:833–43.

50. Ewald JD, Zhou G, Lu Y, Kolic J, Ellis C, Johnson JD, et al. Web-based multi-omics integration using the Analyst software suite. Nat Protoc. 2024;19(5):1467–97.

51. Zhou G, Soufan O, Ewald J, Hancock REW, Basu N, Xia J. NetworkAnalyst 3.0: a visual analytics platform for comprehensive gene expression profiling and meta-analysis. Nucleic Acids Res. 2019;47(W1):W234–W41.

52. Szklarczyk D, Gable AL, Lyon D, Junge A, Wyder S, Huerta-Cepas J, et al. STRING v11: protein-protein association networks with increased coverage, supporting functional discovery in genome-wide experimental datasets. Nucleic Acids Res. 2019;47(D1):D607–D13.

53. Faubert B, Vincent EE, Griss T, Samborska B, Izreig S, Svensson RU, et al. Loss of the tumor suppressor LKB1 promotes metabolic reprogramming of cancer cells via HIF-1α. Proc Natl Acad Sci U S A. 2014;111(7):2554–9.

54. Lu Q, Zong W, Zhang M, Chen Z, Yang Z. The Overlooked Transformation Mechanisms of VLCFAs: Peroxisomal β-Oxidation. Agriculture. 2022;12(7):947.

55. Li N, Li M, Hong W, Shao J, Xu H, Shimano H, et al. Brg1 regulates pro-lipogenic transcription by modulating SREBP activity in hepatocytes. Biochim Biophys Acta Mol Basis Dis. 2018;1864(9 Pt B):2881–9.

56. Zeng PY, Berger SL. LKB1 is recruited to the p21/WAF1 promoter by p53 to mediate transcriptional activation. Cancer Res. 2006;66(22):10701–8.

57. Bungard D, Fuerth BJ, Zeng P-Y, Faubert B, Maas NL, Viollet B, et al. Signaling Kinase AMPK Activates Stress-Promoted Transcription via Histone H2B Phosphorylation. Science. 2010;329(5996):1201–5.

58. Hou X, Liu JE, Liu W, Liu CY, Liu ZY, Sun ZY. A new role of NUAK1: directly phosphorylating p53 and regulating cell proliferation. Oncogene. 2011;30(26):2933–42.

59. Lee SJ, Kang BW, Chae YS, Kim HJ, Park SY, Park JS, et al. Genetic variations in STK11, PRKAA1, and TSC1 associated with prognosis for patients with colorectal cancer. Ann Surg Oncol. 2014;21 Suppl 4:S634–9.

60. Nakatsuka T, Tateishi K, Kudo Y, Yamamoto K, Nakagawa H, Fujiwara H, et al. Impact of histone demethylase KDM3A-dependent AP-1 transactivity on hepatotumorigenesis induced by PI3K activation. Oncogene. 2017;36(45):6262–71.

61. Januario T, Ye X, Bainer R, Alicke B, Smith T, Haley B, et al. PRC2-mediated repression of SMARCA2 predicts EZH2 inhibitor activity in SWI/SNF mutant tumors. Proc Natl Acad Sci U S A. 2017;114(46):12249–54.

62. Kolberg L, Raudvere U, Kuzmin I, Adler P, Vilo J, Peterson H. g:Profiler-interoperable web service for functional enrichment analysis and gene identifier mapping (2023 update). Nucleic Acids Res. 2023;51(W1):W207–w12.

63. Liu P, Ewald J, Pang Z, Legrand E, Jeon YS, Sangiovanni J, et al. ExpressAnalyst: A unified platform for RNA-sequencing analysis in non-model species. Nature Communications. 2023;14(1):2995.

